# Four QTL underlie resistance to a microsporidian parasite that may drive genome evolution in its *Daphnia* host

**DOI:** 10.1101/847194

**Authors:** Devon Keller, Devin Kirk, Pepijn Luijckx

## Abstract

Despite its pivotal role in evolutionary and ecological processes the genetic architecture underlying host-parasite interactions remains understudied. Here we use a quantitative trait loci approach to identify regions in the *Daphnia magna* genome that provide resistance against its microsporidium parasite *Ordospora colligata*. The probability that *Daphnia* became infected was affected by a single locus and an interaction between two additional loci. A fourth locus influenced the number of spores that grew within the host. Comparing our findings to previously published genetic work on *Daphnia magna* revealed that two of these loci may be the same as detected for another microsporidium parasite, suggesting a general immune response to this group of pathogens. More importantly, this comparison revealed that two regions previously identified to be under selection coincided with parasite resistance loci, highlighting the pivotal role parasites may play in shaping the host genome.

## Introduction

Infectious diseases can cause substantial mortality in host populations and thereby affect ecological and evolutionary processes. For example, by increasing mortality of a superior competitor parasites may facilitate species coexistence (Hatcher et al. 2006) or aid invasive species by parasite spill over (Strauss et al. 2012 and references therein). Parasites can also be important selective agents that can shape the genetic structure of host populations (Betts et al. 2018). Indeed, a trematode parasite of New Zealand freshwater snails has been shown to alter the genotype frequencies in its host population by preferentially infecting and reducing fecundity in common snail genotypes (Dybdahl and Lively 1998). The observation that host resistance genes (i.e. genes that prevent or impede parasite establishment or growth within the host) are highly diverse and rapidly evolving further highlights the role parasites have in shaping the genetic architecture of their host (Hammond-Kosack and Jones 1997; Bonneaud et al. 2011; Zhang et al. 2019). However, although ecological and evolutionary theory regularly make assumptions regarding the genetic architecture of parasite resistance, in many cases the actual genetic architecture remains unknown (Ebert 2018). How many genes underlie parasite resistance? How do they interact? And how are they organized in the genome? Addressing these questions is key if we are to better understand the complex ecological and evolutionary processes that parasites are involved in.

Evolutionary and ecological theories have assumed a wide range of genetic models to describe the genetic architecture of host resistance (e.g. Gene For Gene (Thompson and Burdon 1992) and Matching Allele Models (Luijckx et al. 2013) which has led to diverse and sometimes contrasting predictions. For example, under the assumption of quantitative resistance there may be no effect of varying levels of resistance on the occurrence of disease outbreaks (Springbett et al. 2003), while a qualitative genetic assumption suggests large effects (King and Lively 2012). Similarly, genetic architecture plays a key role in the evolutionary debate as to why there is considerable genetic variation for disease-related traits in natural populations despite evolutionary forces that continuously reduce genetic variation (i.e. drift and directional selection). For instance, in models where selection pressure imposed by the parasite on the host follows negative frequency-dependent selection (NFDS, Red Queen models), genetic variation can be maintained indefinitely but only with specific assumptions regarding the genetic architecture (Otto and Nuismer 2004) which have rarely been demonstrated empirically (but see, Metzger et al. 2016). Under NFDS, pathogens adapt to the most common host genotype, which is subsequently outcompeted by a rare host genotype that is resistant to the prevailing pathogen. This continues until this host itself becomes common, and the cycles repeats. To lead to such cycles, the genetic architecture underlying host resistance needs to be based on a small number of loci (five or less (Otto and Nuismer, 2004). In addition, hosts that are resistant to all pathogen genotypes and pathogens that are able to infect all hosts should be absent for NFDS to occur (Luijckx *et al.*, 2013).

A study on the water flea *Daphnia magna* and its microsporidian parasite *Ordospora colligata* (OC) revealed high levels of variation for host resistance and parasite infectivity (Refardt and Ebert 2007, Stanic et al. unpublished) Furthermore, with evidence for highly specialised parasite strains and absence of universally infective parasites (Refardt and Ebert 2007, Stanic et al. unpublished) and evidence for evolution in response to selection by *Ordospora* (Capual and Ebert 2003), this host-parasite system meets the prerequisites for coevolution by negative frequency-dependent selection. The nature of the genetic architecture underlying host specificity, however, remains unknown. Here we use quantitative trait loci (QTL) to identify regions in the host genome that are associated with variation in host resistance against one strain of *Ordospora*. We show that the genetic architecture underlying resistance to this strain is relatively simple, consistent with theory on coevolution by NFDS. Resistance to infection is determined by three QTL that explain 33% of the observed variation, and parasite burden within infected hosts is determined by one QTL which explains 18% of variation. Furthermore, a comparison with previous studies reveals that microsporidian parasites may exert substantial selection on *Daphnia* populations.

## Materials and Methods

### Study system

The water flea *D. magna* inhabits fresh water lakes and ponds across the northern hemisphere where it plays a key role in ecosystem functioning (e.g. by grazing down phytoplankton, influencing water transparency, and preventing the outbreak of algal blooms). Throughout its range it is infected by the microsporidian gut parasite *O. colligata* (Ebert 2005). This parasite shows local adaptation (Refardt and Ebert 2007, Stanic et al. unpublished) and can reach high prevalence (up to 100%) in natural populations (Ebert 2005). Furthermore, there is variation in resistance within populations, with some hosts within the same population being either fully resistant or susceptible (Refardt and Ebert 2007, Stanic et al. unpublished). Infection with *Ordospora* occurs when *Daphnia* inadvertently ingest environmental spores of the parasite while filter feeding. The parasite subsequently invades the epithelial gut cells where it reproduces intracellularly and eventually lyses the cells. After being released, the spores either infect neighbouring gut cells or are released with the faeces into the environment where they can infect new hosts.

### The QTL Panel

The QTL panel used for this study has been previously utilized to identify loci in *D. magna* for other traits, including resistance to several different pathogens. We refer interested readers to Routtu et al (2010) and Routtu et al (2014) for details on the genetic map and the F2 panel. In short, since *D. magna* reproduce via facultative parthenogenesis, genetic crosses can be performed. These crosses result in recombinant offspring that can subsequently be maintained asexually in clonal populations. A standing panel of *D. magna* F2 lines, genotyped at 1324 SNP-markers, is maintained in the laboratory of Dieter Ebert (http://evolution.unibas.ch/ebert/research/qtl/index.htm). This panel was created by selfing a single hybrid F1 clone that was obtained by crossing two parental genotypes that are divergent for numerous life history traits, including parasite resistance. Samples for the parents were originally collected from Finland, Tvärminne (X-clone) and Germany, Munich (I-clone). These genotypes were selfed three times and once, respectively, to obtain the maternal (Xinb3, BB-genotype) and paternal (Iinb1, AA-genotype) genotypes. In total we used 186 different clonal F2 lines from the standing QTL panel to identify resistance loci in *D. magna* against *Ordospora* strain 2, which originated from Belgium.

### Phenotyping

Prior to assessing resistance against *Ordospora*, the *Daphnia* QTL panel was maintained for 8 weeks under standard conditions to minimize maternal effects. Ten to twelve adult females were kept in 400 ml glass microcosms filled with Artificial *Daphnia* Medium (ADaM; Klüttgen et al. 1994, modified to use only 5% of the recommended SiO2 concentration). Animals were transferred to clean microcosms containing fresh medium twice per week and fed three times per week with 125 million batch-cultured green algae (*Monoraphidium minutum*, SAG 278-3, Algae Collection University of Goettingen). Microcosms were kept in a biochamber with a 16:8 light:dark cycle, at 20°C. Phenotyping of *Ordospora* resistance was initiated by collecting six and twelve females juveniles (up to 48 hours old) respectively from each of the F2-lines and both parental lines. Animals were individually placed into separate 100 ml microcosms filled with 80 ml ADaM and 15 million algae. When juveniles were between 3 and 5 days old they were exposed to 50,000 spores of *Ordospora*. The spore solution was prepared by homogenizing a large quantity of infected hosts, and spore concentrations were quantified using a hemocytometer and phase contrast microscopy (400x magnification). Animals were exposed to the parasite for seven days, after which they were transferred to fresh medium. All individuals were fed a diet of 15 million algae three times per week throughout the experiment and were transferred to fresh medium twice per week. The experiment was terminated when animals were 30 days old, at which point all animals were sacrificed and dissected. Dissections and inspection under phase contrast microscopy allowed us to determine infection status (infected or not) as well as quantify the total number of parasite clusters (parasite burden) within the host. Any individuals that died before the end of the experiment were dissected within 24 hours of death, and infection status was recorded.

### Analysis

We measured two types of resistance: resistance to infection (infectivity) and resistance to within-host growth (parasite burden). Infectivity was scored as the proportion of replicates that became infected, excluding any individuals that died within 10 days from the start of the exposure as it is difficult to identify *Ordospora* in the early stages of infection. In addition, we excluded any F2 line that had fewer than three replicates (small sample size was caused by early deaths or undetermined infection status due to dissection failures). Burden was measured as the mean number of spore clusters per F2 line, only including infected replicates from the last day of the experiment (day 30) to avoid interpreting individuals that died early (where the parasite had less time to develop) as more resistant. Burden data was square root transformed to deal with deviations from normality. Analysis for both infectivity and burden were performed using R statistical software version 3.3.1 (R Development Core Team 2016) and QTL were identified using the R package R/qtl version 1.40-8 (Broman et al. 2003). To identify QTL, we performed single genome scans for both traits followed by two dimensional scans to detect epistatic interactions between QTL. Candidate QTL were joined into a single model, and their location was further refined by moving each QTL to the position with the highest likelihood. We used analysis of variance to estimate the proportion of total variance explained by the fitted models. Here, we report the results of the Haley-Knott regression method, which is robust to the use of proportion data. Although this method can be sensitive to epistasis and linkage, the extended Haley-Knott method (which addresses these shortcomings, Feenstra et al. 2006) gave near identical results for single genome scans (results not shown). Genome-wide significance levels for infectivity and burden were calculated using 10,000 permutation tests. For QTLs underlying infection, significant (α < 0.05) and suggested (α < 0.10) QTLs corresponded to LOD scores of 3.79 and 3.47, respectively, while for QTLs underlying spore burden these values corresponded to 3.82 and 3.46.

### Comparison with previous studies

To compare our findings to previous work we obtained the 95% confidence intervals of QTL identified for other pathogens in *D. magna* (Routtu and Ebert 2015; Krebs et al. 2017). As confidence intervals were not always available from the original publications, we reanalysed the original data sets (data kindly provided by D. Ebert) for all QTL with potential overlap with the QTL we identified for *Ordospora*. Two studies on *Hamiltosporidium tvaerminnensis* (formerly known as Octosporia bayeri, Haag et al. 2011) showed potential overlap with the QTL we identified. We reanalysed the QTL identified for *Hamiltosporidium* for spore burden following both horizontal and vertical transmission, and the ability of the parasite to persist in the host population for over 30 weeks (Routtu and Ebert 2015; Krebs et al. 2017). For both studies we used single dimensional scans to obtain the LOD profiles, calculated the 95% confidence intervals and fitted the genetic models specified in their respective publications if multiple QTL were identified on the same linkage group (i.e. chromosome). Finally, we checked if any of the QTL identified overlapped with areas in the *Daphnia* genome found to be under selection (Bourgeois et al. 2017). As this study used a different version of the genome assembly than the previous studies (version 2.4 vs version 2.3) we used BLAST to find the areas under selection and markers associated with the QTL in the newest *D. magna* assembly (Pacbio, Fields and Ebert, unpublished) to verify their positions.

## Results

### Infectivity

The maternal line (Xinb3) was more susceptible to *Ordospora* strain 2 (8 out of 10 replicates infected) than the paternal line (Iinb1, 5 out of 12 infected), although this difference was not significant (P = 0.099 fisher exact test). The F1 hybrid was more resistant than either of the parents with only one replicate out of twelve infected (P = 0.0023). Out of the F2-lines, 30% (n=50) were fully resistant to *Ordospora* strain 2, while in the remaining 70% (n=114) one or more replicates were infected, with only a few lines (4%, n=6) fully susceptible (Fig. 1A). A single-QTL genome scan revealed a QTL for infectivity (scored as the proportion of replicates infected) at the beginning of linkage group 1 with a LOD score of 4.9 explaining ~13% of the observed phenotypic variation (Table 1, Fig. 1B, Table S1). As expected from the parental genotype, individuals carrying alleles from the maternal line (BB or AB) were more susceptible than those that had the paternal genotype (AA, Fig. 2D). Variation explained increased to 17.5% when infectivity was considered a binary trait (i.e. a F2 line was considered susceptible when a single replicate was infected), but due to a lack of convergence in scans for interactions between two QTLs this approach was not further pursued. A two-QTL scan on the proportion of replicates infected did suggest (α < 0.1) the presence of an additional pair of interacting QTL located on linkage group 7 and 8 explaining 23% of the observed phenotypic variation (Fig. 1B, table 1). Individuals carrying the maternal genotype on linkage group 7 (BB, AB) were more susceptible to *Ordospora*, but only if they carried alleles of the paternal genotype (AA, AB) on linkage group 8. Additionally, homozygous individuals (AA) were highly susceptible (~75% of replicates infected) on a BB background (Fig. 1F). A model combining the three QTL and the interaction captured 33% of the phenotypic variation in infectivity (Table 1).

**Table 1:** phenotypic variation explained by the QTL detected for infectivity and spore burden. The full QTL model for infectivity which combined the QTL detected in single dimensional scans (Q1) and QTL detected in two dimensional scans (Q7 and Q8) could explain ~32% of the observed variation. For burden within the host we detected a single QTL (Q6) which explained ~19% of the observed variation. All models were highly significant (P < 0.0001).

| Trait | model | df | Mean-square | LOD | % of variance explained |
| --- | --- | --- | --- | --- | --- |
| Infectivity (n=164) | y~Q1 | 2 | 0.899 | 4.90 | 12.86 |
|  | y~Q7*Q8 | 8 | 0.396 | 9.15 | 22.66 |
|  | y~Q1+Q7*Q8 | 10 | 0.442 | 13.62 | 31.79 |
| Burden (n=114) | y~Q6 | 2 | 648.31 | 5.14 | 18.76 |

**Figure 1:**
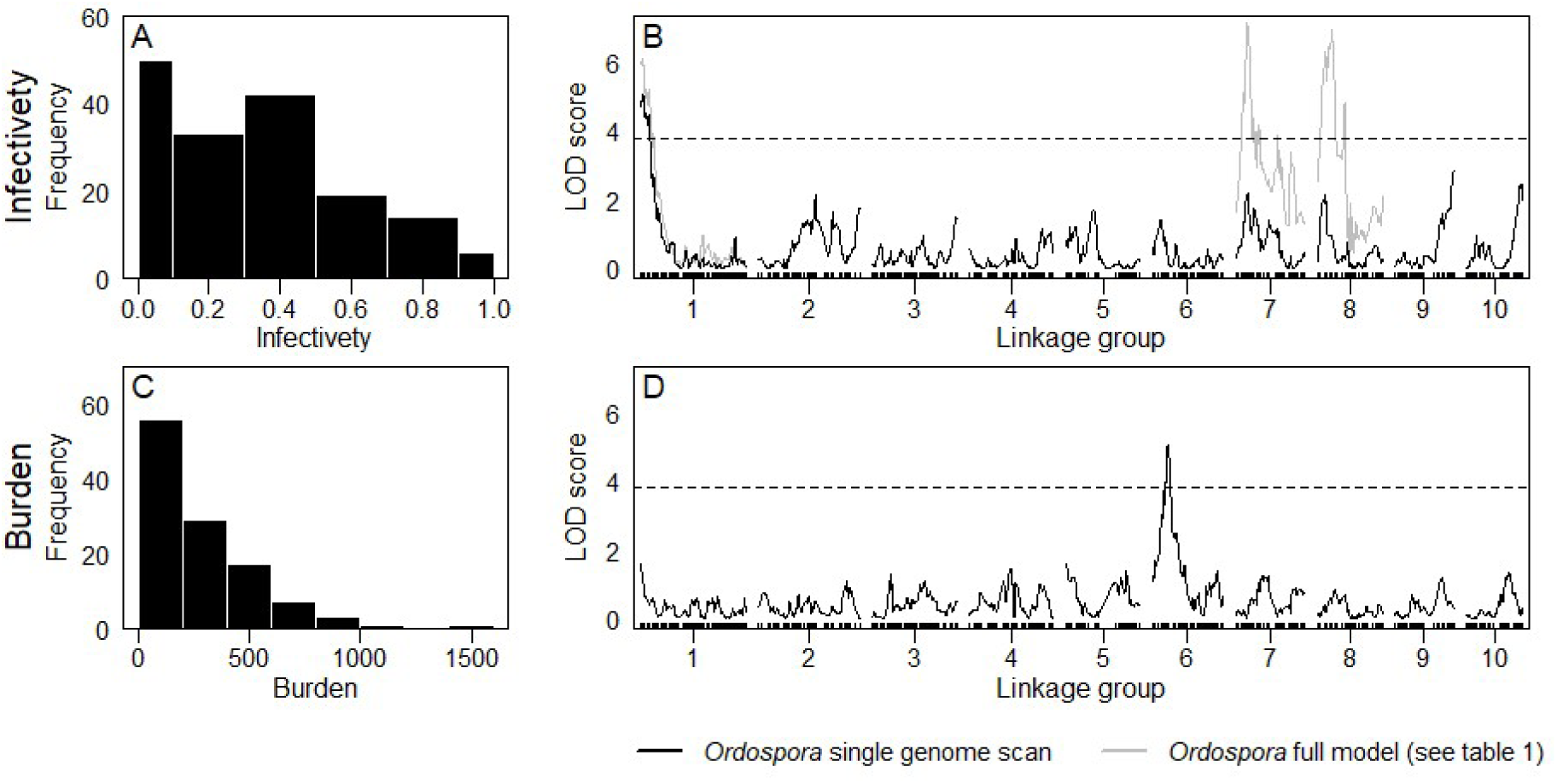
QTL mapping of resistance and burden of the *Daphnia magna* parasite *Ordospora colligata*. **Panel A** shows the frequency distribution of the percentage of replicates that became infected with *Ordospora* for each of the F2 clones. **Panel B** shows the results of the QTL mapping for both the single genome scan for infectivity (black line) and the full QTL model which includes the suggestive interaction (α = 0.1) between QTL on linkage group 7 and 8. **Panel C** shows the frequency distribution of the number of spore clusters within the host and **Panel D** the results of the QTL mapping of the single genome scan for within-host burden. The dotted black line in panels B and D represents the LOD threshold for significance add α = 0.05.

**Figure 2:**
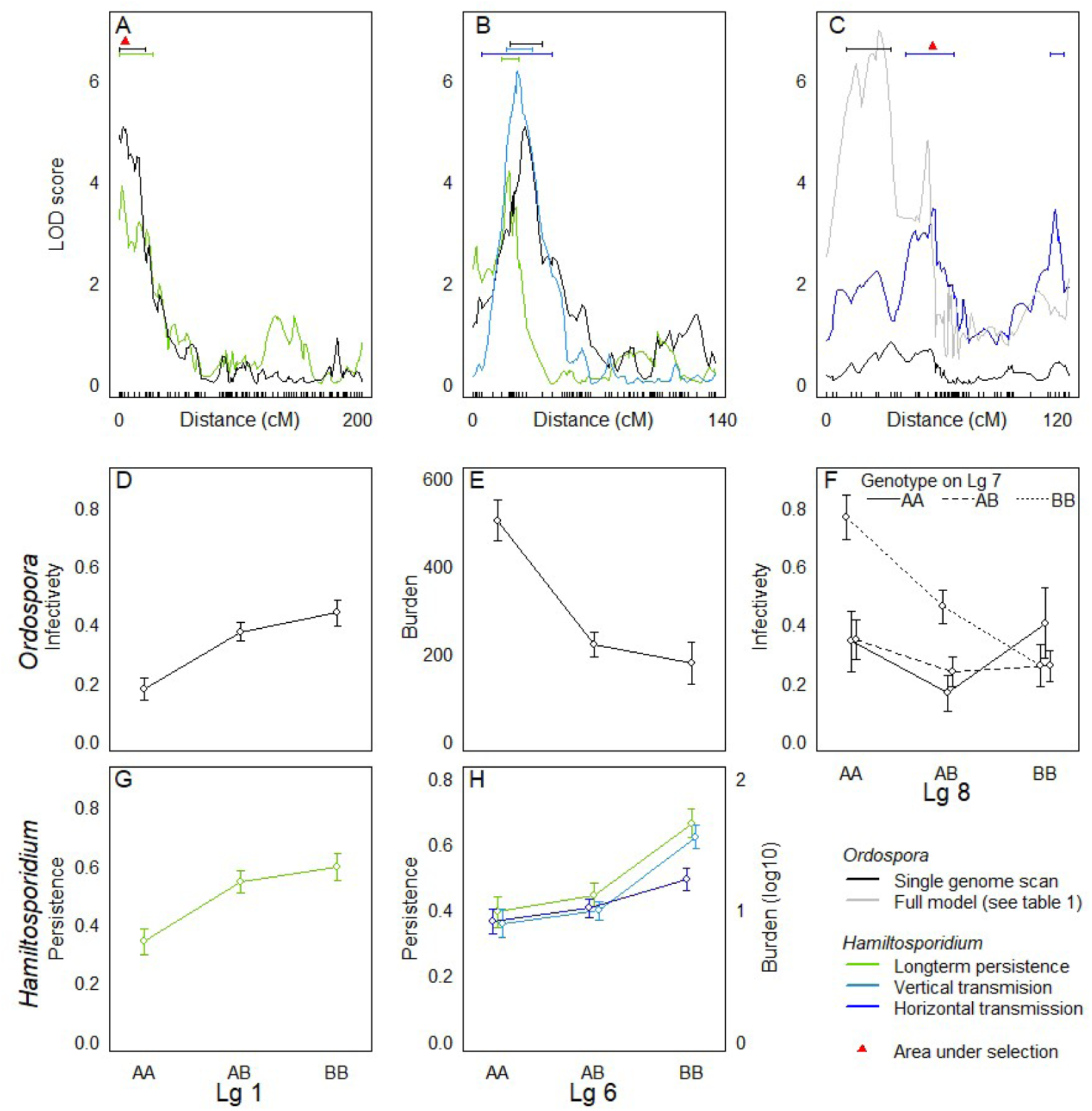
QTL for infectivety and within-host burden of the *Daphnia magna* parasite *Ordospora colligata* in comparison to previously identified resistance QTL for *Hamiltosporidium tvaerminnensis* and areas in the *Daphnia* genome under selection. **Panels A to C** show the QTL profiles of the different studies on linkage groups 1,6 and 8. Ninety-five percent confidence intervals (depicted by the coloured horizontal lines) overlap on linkage group 1 for long-term persistence of *Hamiltosporidium* and *Ordospora* infectivity and occur within an area in the genome under selection (red triangle, **panel A**). The phenotypic effects for both of these QTL are also identical for *Ordospora* and *Hamiltosporidium* (depicted in **panels D** and **G**). Confidence intervals also overlap for QTL on linkage group 6 for *Ordospora* and *Hamiltosporidium*. QTL for horizontal, vertical transmission, and long-term *Hamiltosporidium* persistence coincide with the QTL for within-host burden of *Ordospora* (**panel B**, QTL profile for horizontal transmission, which maps to the same marker as vertical transmission, omitted for clarity). Phenotypic effects in this case are however opposite with individuals homozygous for the “A” allele having a higher *Ordospora* burden (**panel E**) but a lower *Hamiltosporidium* burden (**panel H**). Confidence intervals on linkage group 8 for *Ordospora* resistance and within-host burden for *Hamiltosporidium* did not overlap and these are likely separate loci (**panel C**). Panel F shows the phenotypic effect of the interaction between linkage group 7 and 8 for *Ordospora* resistance.

### Parasite burden

Although both parental lines were susceptible to *Ordospora* strain 2, the infection intensity differed (t_7.5_ = −2.50, P = 0.039), with the maternal line reaching higher numbers of spore clusters (208 spore clusters) after 30 days of exposure than the paternal line (20 spore clusters). The spore numbers in the F1 lines were similar to the paternal line (7 spore clusters). Most lines in the F2 panel (n = 64) had less than 250 spore clusters and only a few (n = 2) reached burdens exceeding 1000 spore clusters (Fig. 1C). A single-QTL genome scan on square root transformed counts of the number of spore clusters revealed a QTL on linkage group 6 with a LOD score of 5.14 which explained 18% of the phenotypic variation in parasite burden in infected hosts (Table 1, Fig. 2D, Table S1). Interestingly, F2 genotypes homozygous for the paternal genotype (AA) carried much higher spore burdens than genotypes with a maternal allele (AB, BB, Fig. 2E), while burden in the parental lines showed the opposite pattern (maternal clone carrying higher burden). A subsequent two-QTL scan did not find any additional significant (α < 0.05) or suggestive (α < 0.1) QTL. The QTL found for infectivity and burden did not overlap and the notion that both types of resistance have an independent genetic basis is reinforced by the absence of a correlation between both traits (supplementary Fig. 1 rho = 0.138, P = 0.155).

### Comparison to other studies

Re-analysis of data from QTL studies on *Hamiltosporidium*, which showed potential overlap with the QTL identified for *Ordospora*, yielded consistent results with the original analysis (Routtu and Ebert 2015; Krebs et al. 2017), with either the same marker identified or a closely linked marker (within 4 cM, Table S1). This analysis also revealed that long-term population-level persistence (30 weeks) of infection with *Hamiltosporidium* was mapped to the same marker that we identified to be associated with infectivity on linkage group 1 for *Ordospora* (Table S1,Fig 2A). Furthermore, the phenotypic effects of both resistance traits are nearly identical (Fig. 2D and 2G) and the same region was previously identified to be under selection (Bourgeois et al. 2017). There was also overlap between a QTL for long-term persistence of *Hamiltosporidium* and the QTL for within-host burden of *Ordospora* on linkage group 6. In addition, the QTL identified by Routtu and Ebert (2015) for spore burden following vertical and horizontal transmission of *Hamiltosporidium* also mapped to the same region (95% confidence intervals of all studies overlapped, Fig. 2B). Interestingly, however, phenotypic effects for *Ordospora* and *Hamiltosporidium* are opposite (e.g. individuals homozygous for the maternal phenotype are susceptible to *Hamiltosporidium* but resistant to *Ordospora*, Fig. 2E and 2H). Routtu and Ebert (2015) also identified two QTL on linkage group 8 but confidence intervals did not overlap with the QTL we identified on this linkage group for infectivity of *Ordospora*. We did, however, find that the marker associated with one of the QTL for *Hamiltosporidium* on linkage group 8 is in proximity (70kb) to another area identified to be under selection (Bourgeois et al. 2017)

## Discussion

Here we identified four QTLs, which together explained a considerable proportion of the variation in host resistance against a microsporidium parasite. Our results provide insight into the genetic architecture of resistance of *Daphnia magna* against *Ordospora colligata*, which underlies local adaptation and the high levels of variation for resistance in this host-parasite system (Refardt and Ebert 2007, Stanic et al. unpublished). Furthermore, the identified QTL operate at different stages of the infection process, highlighting that *Daphnia* has at least two lines of defence against *Ordospora*. Three of the QTL influence how resistant *Daphnia* are to becoming infected (infectivity) while the fourth QTL affects the spore load (burden) of animals that have become infected. A comparison with previously published work (Routtu and Ebert 2015; Bourgeois et al. 2017; Krebs et al. 2017) also revealed that two of the here identified QTL overlap with QTL for resistance against another microsporidian parasite. Finally, this comparison also highlighted the pivotal role parasites may play in shaping the host genome as a QTL for infectivity corresponded to an area previously found to be under selection (Bourgeois et al. 2017).

We observed substantial variation in the F2 panel (186 clonal lines generated by selfing a F1 hybrid) for infectivity, the first line of host defence against *Ordospora*. Our analysis detected one QTL at the beginning of linkage group 1 (Q1) and suggested the presence of a pair of interacting QTL on linkage groups 7 (Q7) and 8 (Q8), which together explained 33% of the observed variation (Fig.1B, Table 1). This is comparable to previous studies, which used the same F2 panel to identify QTL involved in resistance against *Hamiltosporidium tvaerminnensis*, another microsporidian parasite of *Daphnia*. These studies were able to explain 22% and 38% of the observed variation in infection and long term persistence in this related system (J Routtu and Ebert 2015; Krebs, Routtu,s and Ebert 2017). Interestingly, one of the three QTL for long term *Hamiltosporidium* persistence in mesocosm populations identified by Krebs, Routtu and Ebert (2017) was mapped to the same genetic marker on linkage group one as Q1 (Fig 2A, Table S1). This indicates that this QTL may offer simultaneous resistance against both parasites, and potentially microsporidia in general. The fact that this QTL appears to have near identical genotypic effects and dominance in both studies further supports this notion (compare figure 2D and 2G). Both Krebs, Routtu and Ebert (2017) and Routtu and Ebert (2015) identified a QTL on linkage group 6 which influenced spore burden and parasite persistence of *Hamiltosporidium*. Our analysis of the genetics underlying the second line of defence against *Ordospora*, which influences the amount of spore clusters which grow within the host, also found a QTL on linkage group 6 (figure 1D, Table 1). This QTL explained 18% of the phenotypic variation in spore burden with confidence intervals overlapping with those of the previous studies on *Hamiltosporidium* (Fig. 2B). Thus, in addition, to the shared QTL for infectivity on linkage group one, both microsporidia may also share a QTL that influences the spore burden within the host. Notably, however, the effect of the QTL on the spore burden of *Hamiltosporidium* and *Ordospora* is in opposite directions. While the QTL identified for *Hamiltosporidium* makes *Daphnia* individuals that are homozygous for maternal alleles (BB) more susceptible, the QTL identified here for *Ordospora* finds that individuals carrying maternal alleles are more resistant to *Ordospora* (compare Fig. 2E and 2H). This may suggest that either we identified a QTL on a separate locus within the same genomic region as the QTL for *Hamiltospordium*, or that the same locus has opposites effects on these two parasites. Both Routtu and Ebert (2015) and our study also identified QTL on linkage group 8; however, the absence of segregation distortion in our study which was observed for *Hamiltosporidium* and the non-overlapping confidence intervals leads us to believe that these are separate loci (Fig 2C).

We identified separate QTL for infectivity and burden, indicating that both traits act independently to determine *Daphnia* resistance to *Ordospora*. This is further supported by the absence of a correlation between both traits (Fig. S1). It is somewhat surprising that infectivity and spore burden are not correlated to each other, as in our experiment spores released from infected cells are able to re-infect the same individual directly (cell to cell), and one may therefore expect animals with higher resistance to infection to have lower spore loads. One possible explanation is that resistance to infectivity only occurs when the parasite initially invades the gut, and that subsequent within-host spread follows a different mechanism. A potential mechanism could be if *Ordospora* produces separate spores for within- and between-host transmission, as has been suggested for other microsporidia (Dunn, Terry and Smith, 2001; Vizoso and Ebert, 2004; but see Ben-Ami and Urca, 2018 who finds no evidence for seperate functions for different spore types in another microsporidian parasite of *Daphnia*). Independence of different stages in the host defensive cascade have also been demonstrated in other systems (e.g. plants and soil pathogens, Yao and Allen 2006; brood parasites and their avian host, Feeney et al. 2014). Together, these studies support models (Fenton et al. 2012) and calls for subdividing the different stages of the infection process to obtain a more a comprehensive understanding of the epidemiology and evolutionary potential of pathogens (see Hall et al. 2017, for a review of the stepwise infection process). Although we only identified QTL underlying two steps in the host’s process of defending against infection, it is likely that other defensive lines play a role in resisting *Ordospora*. For example, our experimental design did not allow for host migration behaviour, which has been shown to influence parasite resistance (Decaestecker et al. 2002) and has a strong genetic basis (De Meester 1993).

In addition to host QTL potentially affecting different steps of the infection process, there may be additional loci or alleles which act in the defensive steps we identified here. Although the parental genotypes of our study were picked to be as different as possible, the genetic variation captured by the F2 panel represents a subset of the natural variation. Indeed, an explanation for the mismatch between the highly susceptible F2 lines which were homozygous for the paternal genotype that was highly resistant, is that the paternal line was heterozygous for a trait(s) not segregating in the F2 panel (i.e. the F2 panel was based on a single F1 clone which only received half of the genetic material of the paternal line). Results from a pilot experiment also suggest that the genetic architecture of resistance may be more complex, with either more alleles on the loci identified here or additional QTL. This pilot found that three of the seven parasite strains were unable to infect either parent in the QTL panel (Table S2), even though *Daphnia* susceptible to these strains do exist in nature (Refardt and Ebert 2007, Stanic et al. unpublished). Future work testing additional *Ordospora* strains on the QTL panel or using GWAS approaches could help further elucidate the genetic architecture underlying resistance to infection and within-host growth.

Although we lack a detailed understanding of the genetic architecture underlying microsporidia resistance in animals, our finding that resistance to *Ordospora* is coded for by multiple QTL and an epistatic interaction corroborates previous studies. Indeed, as discussed, resistance against the microsporidian *Hamiltosporidium* may even be coded for by the same QTL (Routtu and Ebert 2015; Krebs et al. 2017). Two other studies on the genetic architecture of microsporidia resistance also conclude that multiple QTL underlie resistance, with four QTL identified for resistance in both *C. elegans* and honeybee (*Apis mellifera*) hosts (Huang et al. 2014; Balla et al. 2015). The genetic architecture of microsporidian resistance, with on average 4.2 ± 0.4 QTL and frequent epistasis (80% of cases), is also congruent with the observation that pathogen resistance in other animals is on average coded for by 2.47 ± 1.18 with epistasis detected in 77.4% of studies (see Wilfert and Schmid-Hempel 2008 and references therein). Thus, in general, microsporidian resistance seems to follow the genetic architecture observed for other parasites, but more work is needed to confirm this.

For coevolution to maintain genetic variation via NFDS hosts must show specific resistance to certain parasite strains and parasites must be able to infect specific hosts (Carius et al. 2001). Furthermore, the genetic architecture underlying this specificity needs to be based on few loci (less than five loci; Otto and Nuismer, 2004). Although genotype-genotype interactions have been identified in many host-parasite systems (e.g. snail and trematode, Lively and Dybdahl 2000; mosquito and dengue, Lambrechts 2010; Daphnia and Pasteuria, Luijckx et al. 2011; bumbe bee and Crithidia bombi, Barribeau et al. 2014), information on the genetic architecture underlying genotype – genotype interactions is available for few systems (e.g. Wilfert et al. 2007; Bento et al. 2017). Here, we showed that the genetic architecture of host resistance to *Ordospora* is relatively simple (less than 5 QTL), consistent with the requirements for coevolution by negative frequency-dependent selection. However, as not only the number of QTL but also the interaction (i.e. epistasis) among QTL conferring resistance to different *Ordospora* strains would be critical for the outcome of the evolutionary dynamics, we cannot draw concrete conclusions on the occurrence of NFDS. The absence of parasites able to infect all host types and presence of highly specialised parasite strains (Refardt and Ebert 2007, Stanic et al. unpublished) does suggest that QTL conferring resistance to other *Ordospora* strains may not be independent, or that these QTL carry fitness costs. One potential cost of resistant to *Ordospora* could be susceptibility to *Hamiltosporidium* (i.e. antagonistic pleiotropy), as we find that the same genomic region affects the burden of both microsporidia in opposite directions (both parasites frequently co-occur throughout part of their range (Ebert 2005)). Previous work in another *Daphnia*-pathogen system also identified a genomic region with opposing effects on parasite resistance, depending on the pathogen’s identity. *Daphnia* were either resistant to one strain of the bacterial pathogen *Pasteuria ramosa* or to a different strain, but not resistant to both strains (Luijckx et al. 2013). Modeling work has shown that such a genetic architecture of resistance could support the maintenance of genetic variation at the resistance locus (Luijckx et al. 2013; Engelstädter 2015). Although the locus underlying *Pasteuria* resistance was mapped to linkage group 3 (Bento et al. 2017) and thus does not overlap with the QTL found here, similar dynamics may occur between *Ordospora* and *Hamiltosporidium* if resistance to both microspodia is coded by the same locus.

Regardless of the type of selection (selective sweeps, NFDS or a combination of both), the finding that QTL1 coincided with an area of the genome that was previously found to be under positive selection (Bourgeois et al. 2017) suggests that microsporidia can exert substantial selection pressure on their *Daphnia* host (Fig2A). This is consistent with previous studies on *Ordospora*, which discovered local adaption in natural populations (Refardt and Ebert 2007)(Stanic et al. unpublished), observed changes in genotype frequencies of the host during experimental epidemics (Capual and Ebert 2003), and high prevalence in natural populations (up to 100%, Ebert 2005). Although no genes of known function were located within the direct vicinity of the region under selection, the closest known gene is a Lactosylceramide (Bourgeois et al. 2017) which has been shown to be involved in innate immune processes and is known to bind to different types of pathogens (Zimmerman et al. 1998; Hahn et al. 2003). A comparison of the different genomic studies of *Daphnia* also revealed that an area under selection on linkage group 8 was in proximity (less than 70 kb) to a QTL previously identified to be involved in regulating resistance to *Hamiltosporidium*. This region was also previously associated with the induction of diapause, which is linked to sexual reproduction in *D. magna* (Roulin et al. 2016) and associated with a rhodopsin gene (a photoreceptor) (Bourgeois et al. 2017). In natural populations, induction of diapause may increase resistance to parasites either directly (*Hamiltosporidium* prevalence in resting stages is known to be lower; Vizoso, Lass and Ebert, 2005) or indirectly due to the creation of genetically more diverse offspring by meiosis. Intriguingly, Red Queen models which explore the evolution of sex due to selection by parasites often assume a closely linked pathogen resistance loci and a locus that can modify sexual reproduction (Agrawal 2009). With two regions associated with resistance to microsporidia and an additional two regions previously identified to be under selection by the bacterial pathogen *Pasteuria* (Bourgeois et al. 2017), parasites seem to play a major role in shaping the genome of *Daphnia*. Indeed, out of the six regions under selection that were identified so far, four co-localize with loci involved in parasite resistance.

Pathogens are ubiquitous, and evidence that they play a key role in many ecological and evolutionary processes has been accumulating over the last century (e.g. maintenance of genetic variation, Haldane 1949; mate choice, Hamilton and Zuk 1982). Indeed, in addition to our findings, pathogens have also been identified as a major force of selection in humans with over 100 genes identified to be under selection by a pathogens (Fumagalli et al. 2011), have shown to play a major role in selection of MHC genes (Eizaguirre et al. 2012; Kamiya et al. 2014), and may be responsible for maintaining a disease-resistance polymorphism on the Rpm1 locus of *Arabidopsis thaliana* over millions of years (Stahl et al. 1999). The number of studies that identify genes under selection and link phenotype to genotype is increasing. Further investigation into the role of genetic architecture of host resistance will be key to understanding local adaption in natural populations, the maintenance of genetic variation, disease dynamics and coevolution.

## Acknowledgements

We are grateful to Dieter Ebert and Jürgen Hottinger for providing the *Daphnia* F2 panel, data from previously published work and R-codes for analysis and to Meret J. Halter, Peter Fields and Jarkko Routto for assistance with analysis. We also thank Martin Krkosek for the use of his laboratory and facilities and are grateful to the many undergraduates of University of Toronto that assisted us with this project.

