## Supplementory material for "Four QTL underlie resistance to a microsporidian parasite that may drive genome evolution in its *Daphnia* host"

**Supplementary information.**

Devon Keller[1], Devin Kirk[1,2], Pepijn Luijckx[1,3]

[1] Department of Ecology and Evolutionary Biology, University of Toronto, Toronto, Ontario, Canada, M5S 3G5.

[2] Current address: Department of Biology, Stanford University, Stanford, USA.

[3] School of Natural Sciences, Zoology, Trinity College Dublin, Dublin 2, Ireland

**Supplementary Figures**

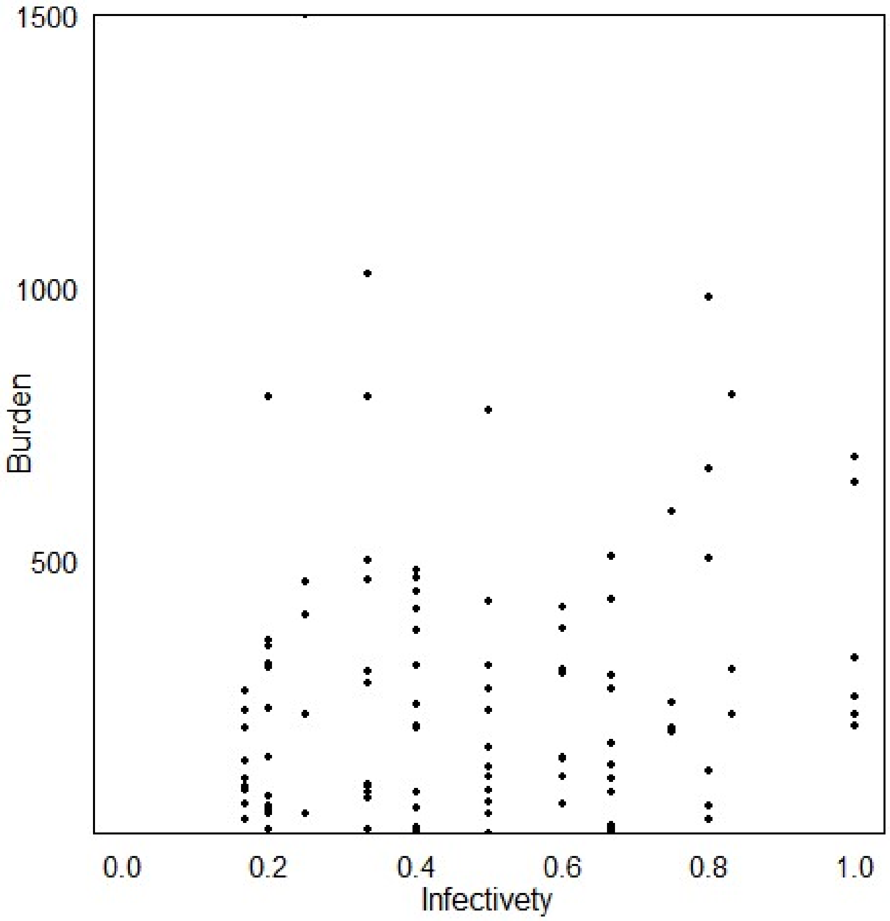


Supplementary Figure 1: Absence of a correlation between infection and burden (Spearman's rank-order correlation, S=175900, p-value = 0.1551).

### Supplementary Tables

**Table S1:** comparison between the here identified loci and other genetic studies pertaining to parasite resistance in *Daphnia magna.* We identified a QTL for infectivity of *Ordospora colligata* at the beginning of linkage group one for which confidence intervals overlapped with the locus that was identified by Krebs et al. 2017 for persistence of the microsporidium *Hamiltosporidium tvaerminnensis* in whole populations. The same region on was also found to be under selection (Bourgeois et al. 2017). Similarly, confidence intervals for QTL of *O. colligata* burden, *H. tvaerminnensis* persistence, vertical and horizontal transmission all overlapped on linkage group 6. However, while QTL were identified for both *O. colligata* and *H. tvaerminnensis* on linkage group 8, confidence intervals for these loci did not overlap. However, we did detect that a region previously found to be under selection on linkage group 8 was within the confidence intervals of a QTL for horizontal transmission of *H. tvaerminnensis.*

| | | | Genome version 2.3 | | | | | | Genome version PacBio | | |
|---|---|---|---|---|---|---|---|---|---|---|---|
| Study | Trait | Type of evidence | Marker | Genome version | Linkage group | Original position (cM) | Reanalysis position (cM) | LOD interval | Contig | Position | Lg |
| Bourgeois* | Area under selection | GWAS | scaffold00568 | *v2.4* | 1 (*1) | **3.7 - 7.3* | - | - | 000016F | ~ 2019389 | 1 |
| Krebs | *Hamiltosporidium,* persistence | QTL | sc01302_1996 | v2.3 | 1 | 2.35 | 2.35 | 1.09 – 25.00 | 000016F | 2199288 | 1 |
| current study | *Ordospora,* infectivity | QTL | sc01302_1996 | v2.3 | 1 | 2.35 | - | 0.00 – 19.00 | 000016F | 2199288 | 1 |
| Routtu** | *Hamiltosporidium,* horizontal transmission | QTL | contig28778_1370 | v2.3 | 6 | 24.01 | 20.51 | 5.10 – 44.48 | 000005F | 1747463 | 8 |
| Routtu** | *Hamiltosporidium,* vertical transmission | QTL | sc00606_2933 | v2.3 | 6 | 24.01 | 25.00 | 19.00 – 33.00 | 000005F | 1747463 | 8 |
| Krebs | *Hamiltosporidium,* persistence | QTL | sc00815_3177 | v2.3 | 6 | 20.34 | 20.34 | 15.97 – 25.43 | 000031F | 1027112 | 8 |
| current study | *Ordospora,* infectivity | QTL | contig28778_1370 | v2.3 | 6 | 27.85 | - | 20.51 - 39.03 | 000005F | 1935968 | 8 |
| current study | *Ordospora,* burden | QTL | sc02738_155 | v2.3 | 7 | 22.00 | - | 18.65 – 28.00 | 000014F | 1463901 | 9 |
| Routtu** | *Hamiltosporidium,* horizontal transmission | QTL | contig14698_81 | v2.3 | 8 | 118.59 | 118.59 | 116.04 – 123 | 000006F | 2796755 | 10 |
| Routtu** | *Hamiltosporidium,* horizontal transmission | QTL | contig29113_349 | v2.3 | 8 | 56.49 | 56.49 | 41.00 – 66.00 | 000013F | 1442280 | 10 |
| Bourgeois* | Area under selection | GWAS* | scaffold01036 | *v2.4* | 8 (*10) | **52.3 - 56.3* | - | - | 000013F | ~ 1513535 | 10 |
| current study | *Ordospora,* infectivity | QTL | sc01110_4114 | v2.3 | 8 | 27 | - | 10 - 32.91 | 000013F | 2591884 | 10 |

* Bourgeois *et al.* 2017 used genome assembly version 2.4, linkage groups given under version 2.3 which was used by the other studies with the 2.4 linkage group between brackets. To determine the approximate area under selection in cM under the 2.3 genome assembly the 2.3 markers flanking the region where located in the newest genome version (PacBio).

** Routtu et al. 2015 used genome assembly version 2.3 and not 2.4 as mentioned in the publication.

**Table S2:** results from a pilot experiment where we exposed multiple asexually propagated individuals of the paternal, maternal and F1 lines to seven strains of *Ordospora colligata.* The F1 was resistant to all strains while the parental and maternal genotypes were substantially infected by one and four strains, respectively.

| | **Ordospora strain** | | | | | | |
|---|---|---|---|---|---|---|---|
| **Host** | OC 2 | OC 3 | OC 5 | OC 6 | OC 7 | OC 8 | OC9 |
| Paternal (*Iinb1*) | 100% (9) | 0% (9) | 0% (12) | 0% (11) | 0% (12) | 0% (12) | 0% (11) |
| Maternal (*Xinb3*) | 100% (10) | 88% (8) | 50% (10) | 91% (11) | 0% (8) | 0% (9) | 13% (8) |
| F1 | 0% (5) | 0% (6) | 0% (6) | 0% (6) | 0% (18) | No data* | 0% (16) |

*Insufficient spores were available to infect the F1 with OC8.
